# Classification of Smartphone Interaction Using Multimodal Physiological Signals with a Brain-Body Spatio-Temporal Transformer

**DOI:** 10.64898/2026.05.03.722573

**Authors:** Prakash Mishra, Vaibhav Kagathara, Tapan K. Gandhi

## Abstract

Distinct smartphone interaction behaviors, like short-form video scrolling and mobile gaming, elicit qualitatively different cognitive and physiological responses. However, such distinctions is often overlooked by approaches that treat smart-phone use as a monolithic behavior. This paper presents Brain-Body Spatio-Temporal Transformer (BB-STT), a unified deep learning framework for classifying interaction-specific physiological signatures from multimodal signals, including EEG, EDA, PPG, and eye-tracking. BB-STT achieves 83.51% accuracy in distinguishing smartphone from non-smartphone activity and 74.13% accuracy in three-class classification of short-form video, gaming, and baseline viewing. The model demonstrates strong generalization with leave-one-subject-out (LOSO) performance that is also comparable to 5-fold cross-validation accuracy. Cross-modal attention emerges as the key component, improving three-class accuracy by 16.74 points through dynamic integration of multimodal signals. Interpretability analysis indicates a hierarchical organization of physiological responses. Eye-tracking features, particularly gaze depth, enable coarse separation between smartphone and non-smartphone activity. In contrast, finer discrimination between passive video viewing and active gaming on smartphones relies on the joint contribution of bilateral pupil dilation and central EEG features. Together, these results demonstrate the potential of multimodal physiological signals for objective, real-time assessment of digital engagement in naturalistic settings.

## I. Introduction

SMARTPHONES have become an integral part of daily life, with over 5.78 billion unique mobile users world-wide and widespread engagement in social media, gaming, and entertainment content [1]. These interactions involve fundamentally distinct cognitive and physiological processes. Short-form video scrolling promotes passive consumption and dopaminergic reward-seeking, inducing low arousal and diminished sustained attention [2], [3], whereas mobile gaming demands strategic planning and fine motor coordination, eliciting high-engagement, flow-like cognitive states [3]. Despite a growing body of evidence linking excessive smartphone use to diminished cognitive performance and psychological distress [4], [5], most existing studies treat smartphone usage as a monolithic behavior, overlooking the qualitatively different physiological engagement profiles induced by different interaction types.

While the ultimate goal of affective computing is to decode subjective emotional experience, a critical prerequisite is to establish that distinct behavioral engagements produce reliably distinguishable physiological signatures. In this work, we address this prerequisite by demonstrating that short-form video scrolling and mobile gaming, behaviors known from prior literature to elicit qualitatively different behavioral engagement patterns [3], [6], produce classifiable multimodal physiological signatures. This provides a foundation for future affect-aware systems incorporating subjective ground truth. Affective and behavioral computing has advanced considerably in emotion recognition from facial expressions, speech, and physiological signals [7], [8], yet the majority of these approaches rely on controlled laboratory stimuli rather than naturalistic, self-directed interactions. Smartphone interactions represent naturalistic, self-directed behaviors that produce ecologically valid physiological responses that more closely approximate real-world behavior [9].

Capturing smartphone engagement behavioral states requires multimodal sensing, as no single modality fully characterizes cognitive and autonomic processes. EEG captures cortical dynamics, while electrodermal activity (EDA), photoplethysmography (PPG), and eye-tracking (ET) contribute complementary peripheral information spanning sympathetic activation, cardiac regulation, and oculomotor engagement [8], [10], [11]. Further, the relevance of each modality varies across behavioral contexts, motivating adaptive fusion strategies rather than fixed combinations.

To address these gaps, this work introduces the Brain-Body Spatio-Temporal Transformer (BB-STT), a multimodal framework integrating multi-scale temporal encoding, spatial attention, and cross-modal attention fusion to dynamically combine neural and peripheral signals. Evaluated across multiple classification tasks, BB-STT achieves strong performance and generalizes across unseen subjects. Interpretability analyses, combining SHapley Additive exPlanations (SHAP) [12], permutation importance, and spatial attention, reveal a two-stage physiological hierarchy, where gaze depth distinguishes device context and EEG and pupillometry resolve interaction-specific engagement.

The main contributions of this work are summarized as follows:

1. **Brain-Body Spatio-Temporal Transformer (BB-STT) for Smartphone Behavioral Engagement Classification**. A unified end-to-end architecture combining multi-scale temporal convolution, transformer-based spatial attention with CLS token readout, and cross-modal attention fusion for classifying physiological signatures of distinct smartphone interaction behaviors. BB-STT achieves 83.51% binary and 74.13% three-class accuracy, outperforming all baselines by up to 19.45 F1 points.
2. **Cross-Modal Attention Fusion for Interaction-Specific Physiological Engagement Profiles**. A cross-modal attention mechanism using EEG as query to dynamically attend to peripheral physiological embeddings, enabling context-sensitive reweighting of autonomic, cardiac, and oculomotor signals. Architectural ablation demonstrates a 16.74-point three-class accuracy improvement over simple concatenation, confirming that modality relevance shifts systematically across behavioral contexts.
3. **Subject-Independent Generalization via LOSO Evaluation**. BB-STT achieves 84.32% LOSO accuracy and 86.73% AUC across 21 participants, marginally exceeding 5-fold CV (83.51%), demonstrating population-level generalization rather than subject-specific memorization, which is a critical requirement for real-world deployment.
4. **A Two-Stage Physiological Hierarchy for Smart-phone Behavioral Decoding**. Through combined SHAP, permutation importance, and spatial attention analysis, we identify that gaze depth dominates device-context separation while EEG occipital gamma and bilateral pupil dilation resolve interaction-type discrimination. This provides a mechanistic interpretation of multimodal contributions to smartphone behavioral state classification.

## II. Related Work

### A. Multimodal EEG-Based Affective and Cognitive State Decoding

The recognition of cognitive and affective states from physiological signals has evolved from unimodal hand-crafted feature pipelines to multimodal deep learning architectures jointly exploiting central and peripheral nervous system recordings. Early convolutional approaches, EEGNet [13] and DeepConvNet [14], established that compact networks can learn discriminative representations from raw EEG, while subsequent reviews [15], [16] documented progression toward deeper architectures for EEG-based affective analysis.

The Transformer [17] marked a paradigm shift in sequential modeling. In the EEG domain, Shi et al. [18] proposed the Multi-band EEG Transformer (MEET), demonstrating that multi-scale spectral decomposition prior to self-attention substantially improves brain-state decoding. Delvigne et al. [19] showed that spatio-temporal self-attention can capture attention-estimation patterns from EEG, and adaptations for tabular [20] and multimodal [21] data further broadened the applicability of attention-based architectures. Graph-based spatial modeling has emerged as a complementary approach: Graph Attention Networks [22] adaptively weight inter-electrode connections, Song et al. [23] pioneered dynamic graph convolutional networks for EEG emotion recognition, and Rahman et al. [24] combined multilayer graph transformers with convolutional networks for functional EEG connectivity analysis.

Cross-modal fusion of EEG with peripheral signals has received increasing attention. Schmidt et al. [8] established the WESAD benchmark for multimodal stress and affect detection. Ding et al. [25] demonstrated that cross-modal attention outperforms early and late fusion for multimodal physiological emotion recognition, directly motivating our fusion design. Zhang et al. [26] introduced DTCA, a dual-branch transformer with cross-attention for EEG and eye-movement fusion, and Jiang et al. [27] proposed ECO-FET leveraging functional brain connectivity and contrastive learning to align EEG and eye-movement representations. Li et al. [28] fused EEG and fNIRS via cross-modality attention for cognitive state decoding, and Zhao et al. [29] proposed multi-query cross-modal attention integrating EEG, ECG, and video for cognitive impairment recognition. In the mobile and wearable space, Yang et al. [30] combined smartphone behavioral and wristband physiological signals for emotion recognition, and Yuvaraj et al. [31] showed EEG and eye gaze fusion achieves 88.56% accuracy for boredom recognition, outperforming any single modality and highlighting the complementary value of neural and oculomotor signals. Despite these advances, to the best of our knowledge, limited prior work has explored multimodal spatio-temporal transformer architectures with cross-modal attention for classifying distinct smartphone interaction behaviors from simultaneously recorded EEG, EDA, PPG, and eye-tracking signals. This gap motivates the proposed approach.

### B. Smartphone Usage and Its Neurophysiological Effects

A substantial evidence base links smartphone use to measurable changes in cognitive performance and affective regulation. At the behavioral level, Hartanto et al. [32] demonstrated through latent variable analysis that executive function deficits are associated with self-reported problematic smartphone use rather than objective screen time, highlighting that engagement quality is more informative than usage quantity. Islambouli et al. [33] showed that purposeless smartphone sessions are associated with regret and reduced well-being, motivating objective engagement monitoring systems.

At the neural level, Ward et al. [34] demonstrated that smartphone presence alone reduces cognitive capacity. Yan et al. [2] linked short-form video consumption to diminished prefrontal theta activity, reflecting reduced executive control relative to other cognitive tasks. Satani et al. [35] reported alpha suppression and sustained beta/gamma activity during social media engagement, consistent with visual cortical arousal and hedonic processing. Pathak et al. [36] identified hedonic motivation and attention regulation difficulties as key neuropsychological drivers of compulsive short-video consumption. At the meta-analytic level, Nguyen et al. [3] confirmed significant negative associations between heavy short-form video use and attention control across 71 studies and approximately 100,000 participants. Collectively, this evidence establishes that different smartphone interaction types produce neurophysiologically distinct engagement states but does not address whether these states can be automatically classified from multimodal physiological recordings in real time.

### C. Smartphone Interaction Classification from Physiological Signals

While the preceding literature establishes that different smartphone engagement types produce distinct neurophysiological signatures, automatic classification of these signatures remains largely unexplored. Hussain et al. [37] applied machine learning to EEG for human activity recognition using explainable AI, and Hafeez et al. [6] developed an EEG framework for classifying gaming expertise, confirming that active play produces distinguishable neural signatures in attention and engagement-related frequency bands. Ciman and Wac [9] demonstrated that smartphone interaction gestures alone carry sufficient information for stress assessment, indicating that behavioral signals during everyday smartphone use encode meaningful engagement content. Huang et al. [38] applied machine learning to EEG functional connectivity for classifying internet addiction patterns, showing that neurophysiological markers can distinguish pathological from healthy digital engagement. These studies, however, focus on binary stress or addiction assessments rather than fine-grained classification of distinct interaction types and their associated physiological engagement profiles.

Passive brain-computer interface (BCI) research provides the conceptual foundation for our approach. Zander and Kothe [39] articulated the vision of passive BCIs continuously monitoring cognitive and affective states without deliberate user intent. The present work realizes a concrete instance of this vision by demonstrating that short-form video scrolling and mobile gaming produce classifiable multimodal physiological engagement signatures decodable in naturalistic settings, without requiring explicit user input or subjective affective labels. To the best of our knowledge, no existing work has used multimodal spatio-temporal information with cross-modal attention to classify distinct smartphone interaction behaviors from simultaneously recorded EEG, EDA, PPG, and ET, constituting the central contribution of this paper. BB-STT leverages multi-scale temporal convolutions for band-specific EEG dynamics, a spatial transformer whose self-attention acts as a data-driven soft adjacency over electrodes, and EEG-queried cross-modal attention to integrate autonomic and oculomotor context. This unified spatio-temporal-multimodal design enables BB-STT to outperform unimodal and fixed-fusion baselines across all classification tasks.

## III. Dataset and Preprocessing

### A. Data Description

Multimodal physiological data were collected from 23 healthy participants (aged 20–32 years; 7 female) with no history of neurological or psychiatric disorders. All participants provided written informed consent prior to participation, and the study protocol received approval from the Institutional Ethics Committee. Two participants (sub-03 and sub-04) were excluded from analysis because their self-selected smartphone activity (reading) did not correspond to either of the two target interaction behaviors under investigation and could not be assigned to either the short-form video or gaming group. The final analysis cohort comprised 21 participants. The complete multimodal dataset is publicly available on OpenNeuro [40].

The experimental paradigm comprised two conditions per participant, presented in counterbalanced order.

#### Smartphone Usage (∼10 min)

Participants were instructed to engage in the smartphone activity they most frequently performed in daily life, without any imposed task constraints. Based on the self-selected activity, participants were assigned to one of two groups representing physiologically distinct interaction behaviors. The Short-Form Video (SV) group (n = 14) engaged in scrolling short-form vertical video content (such as Instagram reels and YouTube shorts), characterized by rapid passive-active content consumption with frequent visual stimulus changes, representing a low-arousal passive consumption behavioral profile. The gaming group (n = 7) played mobile games such as Subway Surfers and Battlefield, requiring sustained visuomotor attention, fine motor coordination, and continuous cognitive engagement, representing a high-arousal active engagement behavioral profile. Group assignment was determined post-hoc from observed behavior rather than imposed by the experimenter, preserving the ecological validity of the interaction.

#### Baseline Viewing (∼5 min)

All participants passively viewed a pre-selected calm, affectively neutral video presented on a separate stationary monitor, serving as the resting behavioral baseline (BL) condition. During all conditions, participants were seated comfortably and instructed to minimize body movements. Recording block boundaries were delineated using stimulus markers embedded directly in the EEG data stream.

### B. Signal Acquisition

Four modalities were recorded simultaneously, capturing complementary central and autonomic nervous system correlates of affective states. A 64-channel EEG was recorded using a Brain Products actiCHamp system with active electrodes arranged according to the extended 10/20 system at a sampling rate of 1000 Hz. Galvanic skin response/electrodermal activity (GSR/EDA), reflecting sympathetic nervous system arousal associated with behavioral engagement, and PPG, providing cardiac dynamics related to cognitive load and autonomic regulation, were recorded via the BrainVision auxiliary GSR and PPG sensors, respectively. Co-sampling at 1000 Hz with the EEG eliminated inter-device synchronization requirements for these modalities. Binocular eye-tracking data were acquired using wearable Tobii Pro Glasses 2 at approximately 100 Hz. Head-mounted inertial measurement unit (IMU) sensors simultaneously recorded 3-axis accelerometer data at ∼ 103 Hz and 3-axis gyroscope data at ∼ 94 Hz. Recorded data streams included gaze direction vectors, pupil center positions, pupil diameter, 2D and 3D gaze points, accelerometer and gyroscope signals, and hardware synchronization markers.

Cross-stream synchronization between EEG and ET was achieved via TTL triggers delivered through the Tobii Pro Glasses 2 synchronization subsystem. At the onset of each recording block, five consecutive TTL triggers were delivered at 1-second intervals to establish a synchronization reference; a single TTL trigger was subsequently delivered every 20 seconds throughout each block to maintain alignment; and five consecutive TTL triggers were delivered at block offset to mark the recording boundary. This protocol ensured subsecond temporal alignment between the EEG and eye-tracking streams across all recording sessions.

### C. Preprocessing

#### 1) Event Segmentation

Recording blocks were delineated using stimulus markers embedded in the EEG stream, identifying smartphone interaction and baseline viewing epochs. Synchronization TTL triggers were used to align EEG and eye-tracking signals to a common time base. Each block was segmented into non-overlapping 1-second windows, yielding a total of 18,900 samples across all participants and conditions.

#### 2) EEG Preprocessing

Raw EEG signals were bandpass filtered (0.5-45 Hz) using a fourth-order zero-phase Butterworth filter to remove slow drifts and high-frequency noise while preserving relevant oscillatory activity. Per-channel z-score normalization was performed using statistics computed within each training fold and applied to the corresponding test fold, preventing data leakage during cross-validation.

#### 3) Peripheral Signal Handling

EDA, PPG, and ET signals were processed using domain-specific feature extraction to account for heterogeneous sampling rates and signal characteristics. For EDA and PPG, 10 features per modality were extracted per segment (Section III-D) and used as inputs to dedicated fully connected feature encoders. ET features (20 dimensions per segment) were similarly pre-extracted due to the event-driven nature of gaze data. This feature-based representation enables the use of established signal processing techniques (e.g., tonic/phasic EDA decomposition and HRV metrics) while maintaining a compact and complementary representation to the raw EEG temporal encoding.

### D. Feature Extraction

Hand-crafted features were extracted from all four modalities for each 1-second segment, yielding 1,576 features per sample (24 × 64 EEG + 20 ET + 10 PPG + 10 EDA) (Table I). These features served dual purposes. Classical machine learning baselines operated on the full 1,576-dimensional vector as a flat input, while deep learning baseline models and BB-STT used the per-channel EEG features (24 per channel) in its feature encoder streams and the peripheral features as inputs to dedicated modality-specific MLP encoders (Section IV). For EEG, 24 features were extracted per channel spanning spectral, temporal, and nonlinear domains. Differential entropy [41] and relative band power [42] were computed across five canonical frequency bands (*δ, θ, α, β, γ*), capturing frequency-specific neural dynamics associated with cognitive and affective engagement [41]. Three Hjorth parameters characterized time-domain signal morphology [43], nine statistical descriptors captured amplitude distribution properties, and spectral and sample entropy [44] quantified signal regularity and complexity.

**TABLE I.**
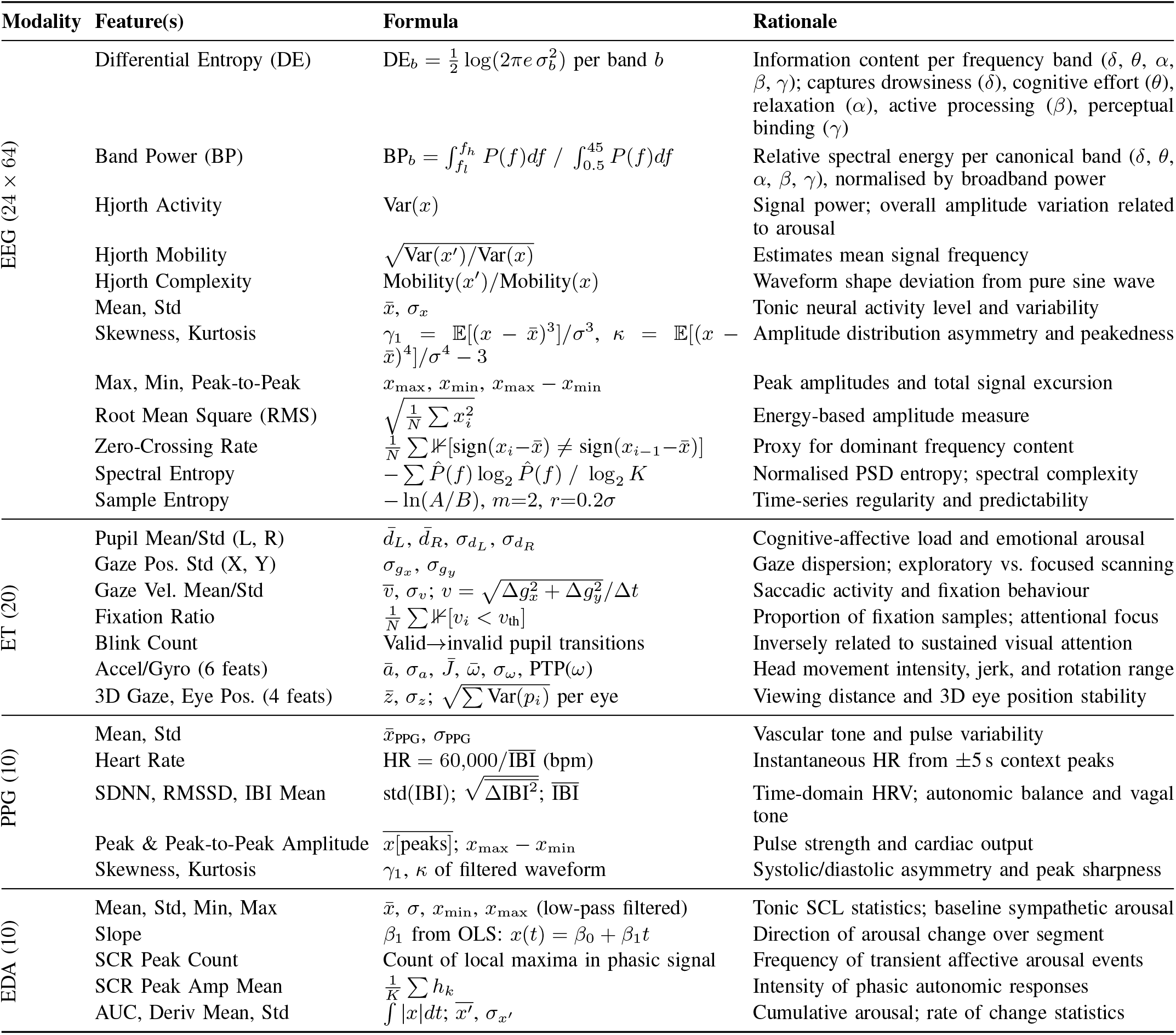
Complete feature set extracted per 1-second segment across all four modalities (24 *×* 64 + 20 + 10 + 10 = 1,576 features total).

Peripheral modality features were designed to capture complementary autonomic and behavioral engagement indices not accessible from EEG alone. EDA features decomposed the electrodermal signal into tonic (skin conductance level (SCL)) and phasic (skin conductance response (SCR)) components, extracting statistical and derivative measures reflecting baseline sympathetic arousal and transient affective reactivity [45], [46]. PPG features included instantaneous heart rate and time-domain heart rate variability (HRV) metrics (SDNN, RMSSD, mean inter-beat intervals (IBI)) derived from detected systolic peaks within a ±5-second context window, providing indices of autonomic regulation and cardiac arousal [10], [47]. Eyetracking features captured pupillometry (mean and standard deviation of bilateral pupil diameter), gaze dynamics (position variability, velocity, fixation ratio, and blink count), head movement from inertial measurement unit sensors, and 3D gaze depth, which collectively encode oculomotor signatures of attentional engagement and cognitive-affective load [11], [48].

### E. Classification Tasks

Five affective-state decoding tasks were defined to provide comprehensive evaluation at different granularities:

#### 1) Binary Classification (Smartphone vs. Baseline)

Distinguishes active smartphone engagement from passive baseline viewing, providing a coarse separation between engaged and neutral affective states.

#### 2) Three-Class Classification (Baseline vs. SV vs. Game)

Extends the binary setting to a more challenging scenario, requiring simultaneous discrimination among baseline (neutral/low arousal), short-form video scrolling (passive engagement), and mobile gaming (active, cognitively demanding engagement).

#### 3) Pairwise Binary Classification

Includes three comparisons, Baseline vs. SV, Baseline vs. Game, and SV vs. Game, to isolate specific physiological contrasts. As reflected in the results, these tasks reveal a hierarchy of discriminability, with SV vs. Game being the most separable and Baseline vs. SV the most challenging.

## IV. Brain-body spatio-temporal transformer

### A. Multi-Scale Temporal Encoder

The temporal encoder processes raw EEG signals **X***∈* ℝ^*C×T*^ (where *C* = 64 channels and *T* = 1000 time samples) through parallel convolutional branches designed to capture oscillatory activity at different frequency scales, each of which carries distinct affective-cognitive information. Four branches with kernel sizes **k** = (250, 125, 50, 25) target frequency bands corresponding approximately to delta (∼4 Hz), theta/alpha (∼ 8 Hz), beta (∼ 20 Hz), and gamma (∼ 40 Hz) bands, respectively [42].

Each branch *b* applies a 1D convolution independently per channel:

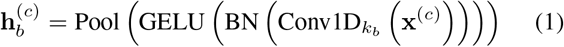

where **x**^(*c*)^ ∈ ℝ^1*×T*^ is the raw signal of channel *c*, 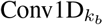 has kernel size *k*_*b*_ with stride *k*_*b*_*/*2, and adaptive average pooling produces a fixed-length representation 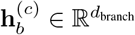 with *d*_branch_ = *d/*4. The outputs of all branches are concatenated and projected:

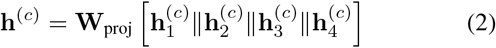

yielding per-channel temporal embeddings **H**_raw_ ∈ ℝ^*C×d*^ where *d* = 64 is the hidden dimension.

### B. Feature Encoder

In parallel with the temporal CNN, the pre-extracted EEG features 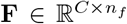 (where *n*_*f*_ = 24 features per channel, see Section III-D) are processed by a two-layer MLP independently per channel:

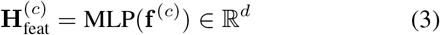

yielding **H**_feat_ ∈ ℝ^*C×d*^. The two streams are fused by concatenation followed by a linear projection:

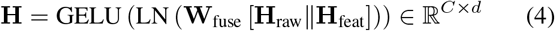

where **W**_fuse_ ∈ ℝ^2*d×d*^. This design allows the model to leverage both learned temporal representations from raw signals and domain-knowledge features (differential entropy, band power, Hjorth parameters) known to carry affective-cognitive information simultaneously.

### C. Spatial Transformer

The spatial transformer models inter-electrode relationships through self-attention, where the attention weights function as a learned soft adjacency matrix over EEG channels [22], [49]. Unlike predefined graph structures based on electrode distances [50], this allows the model to discover affective-state-relevant spatial patterns adaptively.

A learnable class token **z**^[CLS]^ ∈ ℝ^*d*^ is prepended to the channel embeddings, forming the input sequence **Z**_0_ = [**z** ^[CLS]^; **h** ^(1)^; … ; **h** ^(64)^] ∈ ℝ ^65×*d*^.

The sequence is processed by *L* = 2 transformer encoder layers [17], each comprising multi-head self-attention (MHSA) with *K* = 4 heads and a position-wise feed-forward network (FFN):

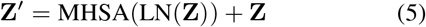

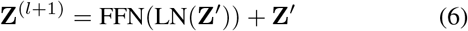

Each attention head *k* computes:

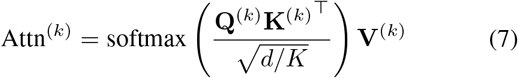

The [CLS] token at the final layer output aggregates information from all channels, yielding the EEG embedding **z**^EEG^ ∈ ℝ^*d*^. The attention maps from the spatial transformer can be visualized as learned brain connectivity graphs reflecting affective-state-specific neural network configurations, providing interpretability.

### D. Peripheral Affective Signal Encoders

#### EDA and PPG Encoders

Pre-extracted EDA features **x**^EDA^ ∈ ℝ^10^, capturing sympathetic arousal correlates of affective engagement, and PPG features **x**^PPG^ ∈ ℝ^10^, capturing cardiac dynamics related to emotional arousal (Section III-D), are each encoded by a single-layer MLP:

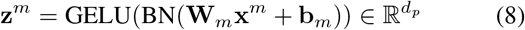

where *m* ∈ {EDA, PPG} and *d*_*p*_ = 64. This design leverages the domain-specific feature extraction (tonic/phasic EDA decomposition for arousal characterization, HRV metrics from PPG for affective autonomic regulation) while projecting the heterogeneous peripheral representations into a common affective embedding space.

#### Eye-Tracking Encoder

Pre-extracted ET features **x**^ET^ ∈ ℝ^20^, capturing oculomotor signatures of attentional engagement and affective processing, are encoded by a single-layer MLP:

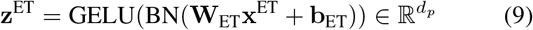

### E. Cross-Modal Affective Attention Fusion

The peripheral affective embeddings {**z**^EDA^, **z**^PPG^, **z**^ET^} are stacked into a matrix 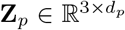 and projected to the EEG embedding dimension via 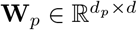. The EEG embedding serves as the query and the peripheral affective embeddings serve as keys and values in a cross-attention mechanism:

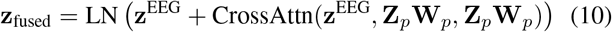

This design intentionally uses EEG as the query modality, allowing the model to selectively attend to complementary peripheral affective information, such as autonomic arousal from EDA and cardiac responses from PPG, while maintaining the EEG representation of central nervous system affective processing as the primary signal [25]. The cross-attention uses 2 heads with a residual connection.

### F. Affective-State Classification Head

The fused multimodal affective representation is passed to a two-layer MLP:

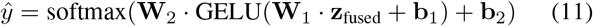

where **W**_1_ ∈ ℝ^*d×d*^, 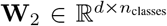 , with dropout between layers. The model is trained end-to-end using cross-entropy loss with class weighting to handle label imbalance.

#### Algorithm 1

BB-STT: Training and Inference Pipeline

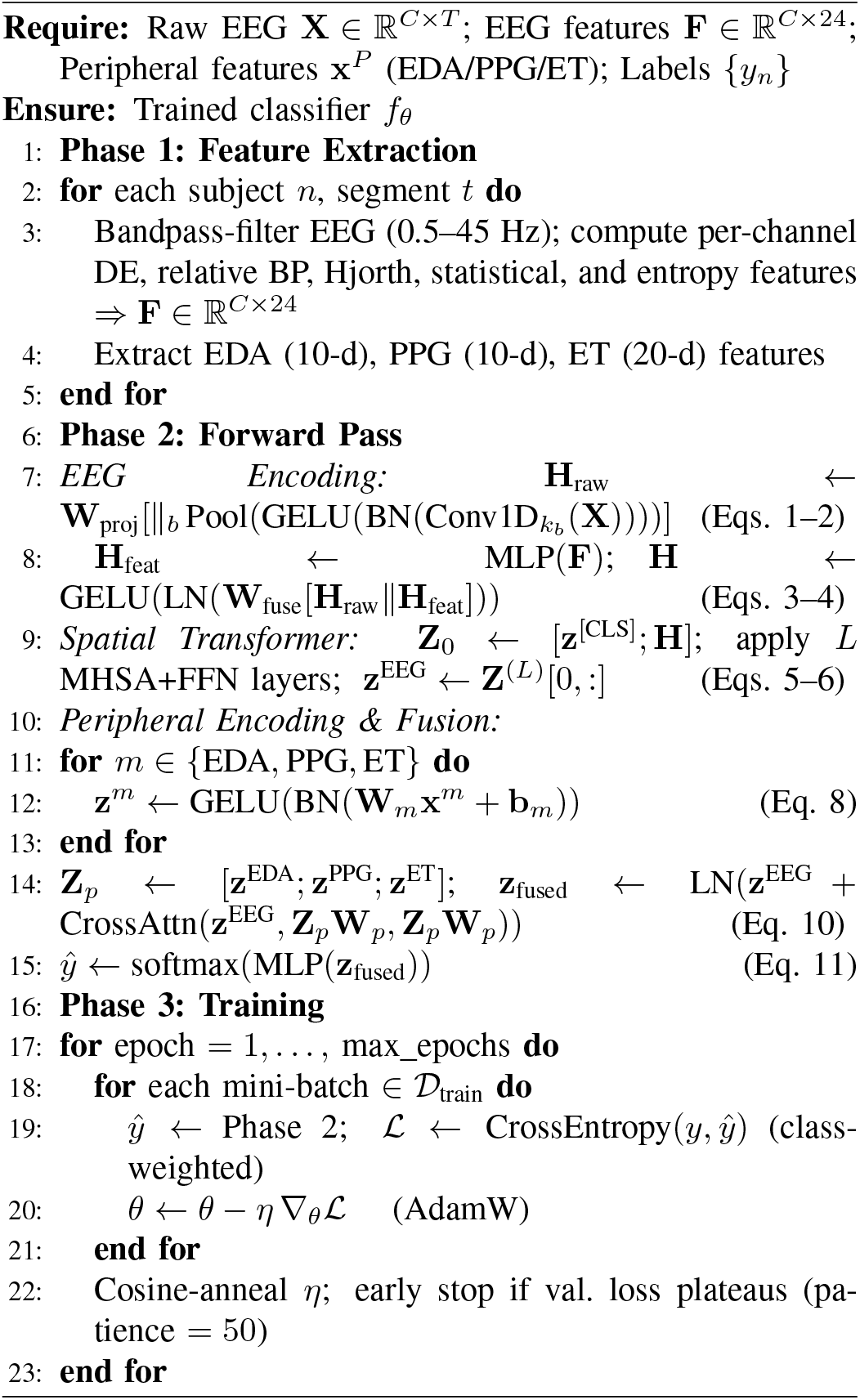

### G. Training Details

All models were implemented in PyTorch and trained on an NVIDIA RTX A4000 GPU with automatic mixed precision (AMP). Key hyperparameters: hidden dimension *d* = 64, spatial transformer layers *L* = 2, attention heads *K* = 4, temporal kernels = (250, 125, 50, 25), peripheral encoder dimension *d*_*p*_ = 64, EEG feature dimension *n*_*f*_ = 24, dropout = 0.3, learning rate = 10^*−*3^ with cosine annealing, batch size = 1024, maximum epochs = 100 with early stopping (patience = 50). The AdamW optimizer with weight decay = 10^*−*4^ was used. Throughout the architecture, LN denotes layer normalisation, BN denotes batch normalisation, and CrossAttn denotes cross-attention (see Section IV-E). The complete training and inference pipeline is summarised in Algorithm 1.

## V. Results

### A. Experimental Setup and Evaluation Protocol

All primary experiments use subject-wise stratified 5-fold cross-validation, in which entire participants are assigned to folds rather than individual segments, preventing data leakage from subject-specific physiological patterns appearing in both training and test sets. At least one gaming subject is included in each test fold to preserve class representation across folds. Each fold contains between 2,700 and 4,700 test segments out of 18,900 total segments. All preprocessing statistics, such as per-channel z-score normalization for EEG and feature scaling for peripheral modalities, are computed exclusively within each training fold and applied to the corresponding held-out test fold.

To assess generalization to unseen individuals, we additionally evaluate BB-STT under leave-one-subject-out (LOSO) cross-validation for the binary task, training on all segments from 20 subjects and testing on the single held-out subject, repeated for all 21 participants. Performance is reported as mean ± standard deviation across folds using accuracy, macro-averaged precision, macro-averaged recall, and macro-averaged F1-score. Macro averaging treats all classes equally regardless of frequency, making it the appropriate primary metric given the 2:1 imbalance between SV and gaming subjects. BB-STT is benchmarked against six classical machine learning baselines operating on the full 1,576-dimensional hand-crafted feature vector and six deep learning baselines trained on the same multimodal inputs as BB-STT.

To characterise the discriminative contribution of individual features, we employ two complementary interpretability methods. Permutation importance (ΔAccuracy) measures each feature’s causal contribution by randomly shuffling its values across test samples and measuring the resulting drop in classification accuracy; a large ΔAccuracy indicates that the feature carries individually irreplaceable information. SHAP (SHapley Additive exPlanations) [12] values, grounded in cooperative game theory, quantify each feature’s average marginal contribution to the model’s output across all possible feature coalitions, capturing both direct effects and interactions with other features. We report mean absolute SHAP values (mean |SHAP|) averaged across test samples. Plotting ΔAccuracy (x-axis) against mean |SHAP| (y-axis) for each feature produces a diagnostic scatter (Fig. 3) that disambiguates two modes of importance: features in the top-right quadrant are both individually irreplaceable (high ΔAccuracy) and consistently influential across feature coalitions (high SHAP), whereas features with high SHAP but low ΔAccuracy are influential yet redundant, their information is shared with correlated features that can compensate when the feature is removed.

### B. Separating Smartphone from Non-Smartphone Engagement

The first and most fundamental question is whether physiological signals recorded during any form of active smartphone use are reliably distinguishable from those recorded during passive baseline monitor viewing. This binary task provides a prerequisite validation step, ensuring that coarse-level separability exists before attempting finer discrimination between specific interaction types. We evaluate this question under two complementary protocols, stratified 5-fold cross-validation providing a within-population estimate (Table II), and leave-one-subject-out (LOSO) cross-validation (Table III) providing a conservative estimate of generalization to previously unseen individuals.

**TABLE II.** Binary affective-state decoding: smartphone vs. baseline (5-fold cv, mean *±* std). best results in bold.

| Type | Model | Accuracy | Precision | Recall | F1-Score |
| --- | --- | --- | --- | --- | --- |
| Classical | LR | 0.6254±0.09 | 0.6055±0.06 | 0.6052±0.05 | 0.5922±0.05 |
| | SVM (RBF) | 0.6757±0.02 | 0.5796±0.10 | 0.5197±0.07 | 0.4460±0.07 |
| | RF | 0.7201±0.09 | 0.7351±0.06 | 0.7112±0.05 | 0.6903±0.05 |
| | XGBoost | 0.6944±0.11 | 0.6977±0.05 | 0.6856±0.04 | 0.6609±0.05 |
| | LightGBM | 0.7039±0.07 | 0.7027±0.03 | 0.7116±0.03 | 0.6835±0.06 |
| | KNN | 0.5986±0.05 | 0.5375±0.10 | 0.5373±0.07 | 0.5335±0.07 |
| DL | MLP | 0.6847±0.02 | 0.4911±0.20 | 0.5439±0.05 | 0.4815±0.10 |
| | EEGNet | 0.6661±0.07 | 0.6163±0.18 | 0.5784±0.08 | 0.5402±0.10 |
| | CNN-1D | 0.6934±0.04 | 0.5235±0.23 | 0.5456±0.07 | 0.4789±0.11 |
| | LSTM | 0.6824±0.03 | 0.4390±0.21 | 0.5237±0.05 | 0.4422±0.08 |
| | CNN-LSTM | 0.7016±0.04 | 0.5295±0.24 | 0.5562±0.07 | 0.4975±0.12 |
| | Transformer | 0.6831±0.02 | 0.5612±0.20 | 0.5337±0.03 | 0.4740±0.06 |
| Ours | **BB-STT** | **0.8351±0.06** | **0.8524±0.05** | **0.7605±0.06** | **0.7835±0.09** |

**TABLE III.** Per-subject loso binary classification results (smartphone vs. baseline). each fold represents one held-out subject.

| Subject | Activity | Acc | Prec | Rec | F1 | AUC |
| --- | --- | --- | --- | --- | --- | --- |
| sub-01 | short-video | 0.8533 | 0.8329 | 0.8442 | 0.8379 | 0.8961 |
| sub-02 | game | 0.8578 | 0.8387 | 0.8683 | 0.8477 | 0.9279 |
| sub-05 | short-video | 0.9011 | 0.9301 | 0.8542 | 0.8792 | 0.9450 |
| sub-06 | short-video | 0.7567 | 0.7363 | 0.7592 | 0.7411 | 0.8531 |
| sub-07 | game | 0.9133 | 0.9346 | 0.8742 | 0.8959 | 0.9330 |
| sub-08 | game | 0.8289 | 0.8979 | 0.7433 | 0.7705 | 0.8353 |
| sub-09 | short-video | 0.8133 | 0.8778 | 0.7233 | 0.7473 | 0.9518 |
| sub-10 | short-video | 0.9078 | 0.9295 | 0.8667 | 0.8889 | 0.9438 |
| sub-11 | game | 0.9244 | 0.9491 | 0.8867 | 0.9093 | 0.9571 |
| sub-12 | short-video | 0.9378 | 0.9340 | 0.9250 | 0.9293 | 0.9533 |
| sub-13 | short-video | 0.7689 | 0.7410 | 0.7258 | 0.7320 | 0.8329 |
| sub-14 | short-video | 0.8967 | 0.9004 | 0.8642 | 0.8786 | 0.9184 |
| sub-15 | short-video | 0.7044 | 0.8328 | 0.5575 | 0.5133 | 0.4305 |
| sub-16 | game | 0.8944 | 0.9134 | 0.8508 | 0.8726 | 0.9574 |
| sub-17 | short-video | 0.8689 | 0.8513 | 0.8558 | 0.8535 | 0.9051 |
| sub-18 | game | 0.7667 | 0.8704 | 0.6500 | 0.6563 | 0.9605 |
| sub-19 | short-video | 0.6656 | 0.3331 | 0.4992 | 0.3996 | 0.4349 |
| sub-20 | game | 0.8222 | 0.8651 | 0.7425 | 0.7670 | 0.7919 |
| sub-21 | short-video | 0.9244 | 0.9476 | 0.8875 | 0.9095 | 0.9678 |
| sub-22 | short-video | 0.7556 | 0.8600 | 0.6342 | 0.6345 | 0.8308 |
| sub-23 | short-video | 0.9456 | 0.9397 | 0.9375 | 0.9386 | 0.9876 |
| **Average -** |  | **0.8432** | **0.8531** | **0.7881** | **0.7906** | **0.8673** |

#### 1) 5-Fold Cross-Validation Results

Among classical models, Random Forest performs best (72.01% accuracy, 69.03% F1), followed by LightGBM and XGBoost, while linear regression and KNN show weaker performance. The consistent gap between accuracy and F1-score indicates limited handling of class imbalance using hand-crafted features alone. Among deep learning baselines, CNN-LSTM achieves the highest accuracy (70.16%), but all models exhibit relatively low F1-scores, suggesting difficulty in learning balanced representations without effective multimodal fusion. EEGNet further highlights the limitation of unimodal approaches (54.02% F1).

In contrast, BB-STT achieves 83.51% accuracy and 78.35% F1-score, outperforming all baselines by a substantial margin, demonstrating the effectiveness of cross-modal attention for multimodal physiological integration.

#### 2) Generalization to Unseen Subjects: LOSO Evaluation

To establish that these learned physiological signatures transfer to individuals absent from training, a prerequisite for any real-world deployment scenario, BB-STT is additionally evaluated under leave-one-subject-out cross-validation across all 21 participants. Across all subjects, BB-STT achieves an average LOSO accuracy of 84.32%, precision of 85.31%, recall of 78.81%, F1-score of 79.06%, and AUC of 86.73%. Crucially, LOSO accuracy marginally exceeds the 5-fold CV result (83.51%), demonstrating that BB-STT learns population-level physiological engagement signatures rather than subject-specific idiosyncrasies. This close correspondence between the two evaluation protocols validates the subject-wise fold assignment strategy and provides strong evidence that the within-population 5-fold estimates are not inflated by within-subject segment similarity. Per-subject performance varies considerably, ranging from 66.56% accuracy (sub-19, short-video) to 94.56% accuracy (sub-23, short-video), with AUC spanning 43.05 to 98.76, which shows individual differences in resting-state EEG, autonomic baseline reactivity, and oculomotor behavior produce subject-specific offsets that a population-trained model must bridge without adaptation.

Taken together, the 5-fold and LOSO results establish that smartphone engagement is both accurately classifiable within a known population and robustly generalizable to new users, providing a strong empirical foundation for the finer-grained behavioral discriminations evaluated in the following sections.

### C. Three-Class Behavioral Discrimination: Baseline, Short-Form Video, and Gaming

Having established binary separability and cross-subject generalization, we evaluate simultaneous three-class discrimination, baseline vs. short-form video vs. gaming, which requires the model to resolve qualitative differences between two forms of smartphone engagement while also distinguishing both from passive viewing (Table IV).

**TABLE IV.** Three-class affective-state decoding: baseline vs. sv vs. game (5-fold cv, mean *±* std). best results in bold.

| Type | Model | Accuracy | Precision | Recall | F1-Score |
| --- | --- | --- | --- | --- | --- |
| Classical | LR | 0.4212±0.10 | 0.4736±0.06 | 0.4228±0.06 | 0.4199±0.04 |
| | SVM (RBF) | 0.4467±0.07 | 0.1489±0.06 | 0.3333±0.05 | 0.2048±0.03 |
| | RF | 0.4179±0.06 | 0.3698±0.06 | 0.3914±0.03 | 0.3100±0.04 |
| | XGBoost | 0.4375±0.10 | 0.4137±0.07 | 0.4293±0.07 | 0.3960±0.07 |
| | LightGBM | 0.4271±0.08 | 0.4566±0.07 | 0.4046±0.05 | 0.3693±0.05 |
| | KNN | 0.3759±0.06 | 0.3752±0.09 | 0.3923±0.08 | 0.3586±0.09 |
| DL | MLP | 0.5264±0.07 | 0.4732±0.22 | 0.5007±0.14 | 0.4475±0.16 |
| | EEGNet | 0.5399±0.14 | 0.4565±0.20 | 0.5730±0.13 | 0.4591±0.16 |
| | CNN-1D | 0.5177±0.07 | 0.5440±0.13 | 0.4686±0.14 | 0.4102±0.14 |
| | LSTM | 0.5574±0.06 | 0.4002±0.16 | 0.4952±0.11 | 0.3854±0.10 |
| | CNN-LSTM | 0.5978±0.06 | 0.5747±0.15 | 0.6146±0.07 | 0.5012±0.06 |
| | Transformer | 0.5560±0.06 | 0.3407±0.08 | 0.4940±0.12 | 0.3838±0.10 |
| Ours | **BB-STT** | **0.7413±0.11** | **0.6566±0.15** | **0.5741±0.09** | **0.6957±0.14** |

As expected, a uniform decrease in performance relative to the binary task is observed across all models. Classical models perform near chance level, with the best classical accuracy at only 44.67% (SVM-RBF) against a three-class chance baseline of 33.3%. LR achieves 41.99% F1 while RF and KNN perform weakest at 31.00% and 35.86% F1 respectively, indicating that the three-class physiological feature space is not linearly or distance-metrically separable with hand-crafted features alone. Deep learning baselines show more substantial improvement, with CNN-LSTM reaching 59.78% accuracy and 50.12% F1 as the strongest baseline, though the persistent gap between accuracy and F1 across all deep baselines reflects residual difficulty in resolving the SV vs. gaming distinction without dedicated multimodal fusion.

BB-STT achieves 74.13% accuracy, 65.66% precision, 57.41% recall, and 69.57% F1-score, outperforming the best baseline by 14.35 percentage points in accuracy and 19.45 points in F1. This substantial gain highlights the effectiveness of cross-modal attention in resolving the challenging SV vs. gaming distinction, where subtle differences require dynamic integration of neural and peripheral physiological signals.

### D. Pairwise Analysis: What Makes Each Condition Physiologically Distinct

Table V presents pairwise classification results isolating the physiological contrast between each condition pair. The three comparisons reveal a clear hierarchy of discriminability and provide interpretable insight into the physiological mechanisms differentiating each behavioral state.

**TABLE V.**
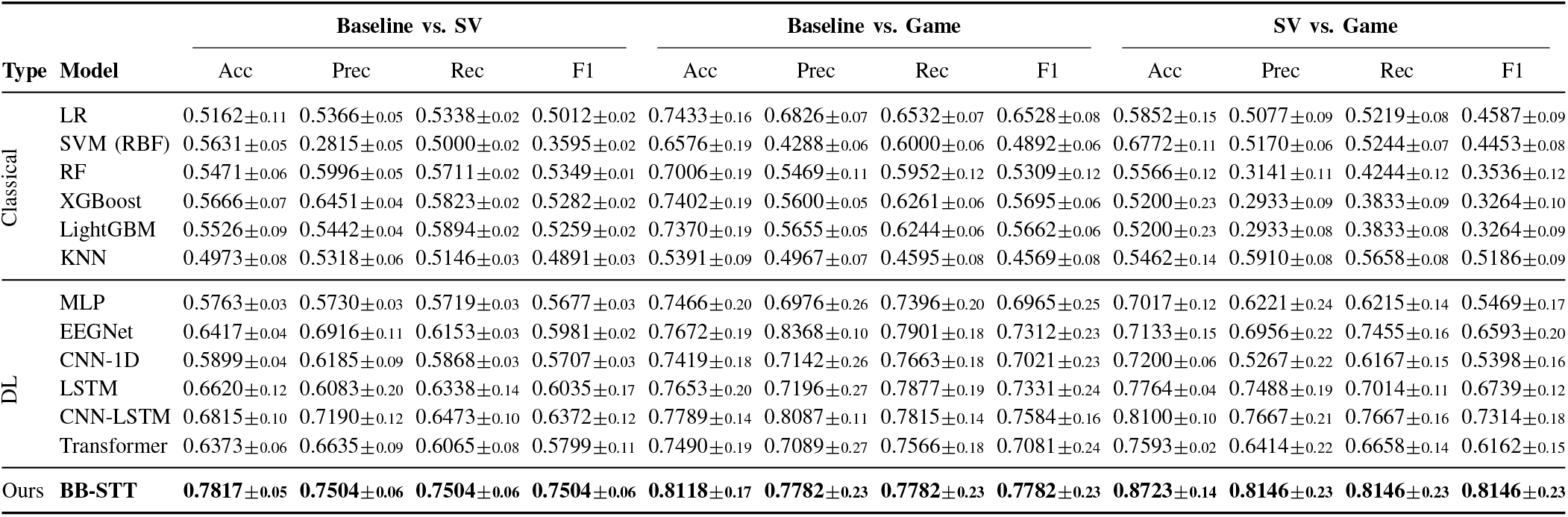
Pairwise affective-state decoding results (5-fold cv, mean *±* std). best results in bold.

| Type | Model | Baseline vs. SV |  |  |  | Baseline vs. Game |  |  |  | SV vs. Game |  |  |  |
| --- | --- | --- | --- | --- | --- | --- | --- | --- | --- | --- | --- | --- | --- |
|  |  | Acc | Prec | Rec | F1 | Acc | Prec | Rec | F1 | Acc | Prec | Rec | F1 |
| Classical | LR | 0.5162±0.11 | 0.5366±0.05 | 0.5338±0.02 | 0.5012±0.02 | 0.7433±0.16 | 0.6826±0.07 | 0.6532±0.07 | 0.6528±0.08 | 0.5852±0.15 | 0.5077±0.09 | 0.5219±0.08 | 0.4587±0.09 |
| | SVM (RBF) | 0.5631±0.05 | 0.2815±0.05 | 0.5000±0.02 | 0.3595±0.02 | 0.6576±0.19 | 0.4288±0.06 | 0.6000±0.06 | 0.4892±0.06 | 0.6772±0.11 | 0.5170±0.06 | 0.5244±0.07 | 0.4453±0.08 |
| | RF | 0.5471±0.06 | 0.5996±0.05 | 0.5711±0.02 | 0.5349±0.01 | 0.7006±0.19 | 0.5469±0.11 | 0.5952±0.12 | 0.5309±0.12 | 0.5566±0.12 | 0.3141±0.11 | 0.4244±0.12 | 0.3536±0.12 |
| | XGBoost | 0.5666±0.07 | 0.6451±0.04 | 0.5823±0.02 | 0.5282±0.02 | 0.7402±0.19 | 0.5600±0.05 | 0.6261±0.06 | 0.5695±0.06 | 0.5200±0.23 | 0.2933±0.09 | 0.3833±0.09 | 0.3264±0.10 |
| | LightGBM | 0.5526±0.09 | 0.5442±0.04 | 0.5894±0.02 | 0.5259±0.02 | 0.7370±0.19 | 0.5655±0.05 | 0.6244±0.06 | 0.5662±0.06 | 0.5200±0.23 | 0.2933±0.08 | 0.3833±0.08 | 0.3264±0.09 |
| | KNN | 0.4973±0.08 | 0.5318±0.06 | 0.5146±0.03 | 0.4891±0.03 | 0.5391±0.09 | 0.4967±0.07 | 0.4595±0.08 | 0.4569±0.08 | 0.5462±0.14 | 0.5910±0.08 | 0.5658±0.08 | 0.5186±0.09 |
| DL | MLP | 0.5763±0.03 | 0.5730±0.03 | 0.5719±0.03 | 0.5677±0.03 | 0.7466±0.20 | 0.6976±0.26 | 0.7396±0.20 | 0.6965±0.25 | 0.7017±0.12 | 0.6221±0.24 | 0.6215±0.14 | 0.5469±0.17 |
| | EEGNet | 0.6417±0.04 | 0.6916±0.11 | 0.6153±0.03 | 0.5981±0.02 | 0.7672±0.19 | 0.8368±0.10 | 0.7901±0.18 | 0.7312±0.23 | 0.7133±0.15 | 0.6956±0.22 | 0.7455±0.16 | 0.6593±0.20 |
| | CNN-1D | 0.5899±0.04 | 0.6185±0.09 | 0.5868±0.03 | 0.5707±0.03 | 0.7419±0.18 | 0.7142±0.26 | 0.7663±0.18 | 0.7021±0.23 | 0.7200±0.06 | 0.5267±0.22 | 0.6167±0.15 | 0.5398±0.16 |
| | LSTM | 0.6620±0.12 | 0.6083±0.20 | 0.6338±0.14 | 0.6035±0.17 | 0.7653±0.20 | 0.7196±0.27 | 0.7877±0.19 | 0.7331±0.24 | 0.7764±0.04 | 0.7488±0.19 | 0.7014±0.11 | 0.6739±0.12 |
| | CNN-LSTM | 0.6815±0.10 | 0.7190±0.12 | 0.6473±0.10 | 0.6372±0.12 | 0.7789±0.14 | 0.8087±0.11 | 0.7815±0.14 | 0.7584±0.16 | 0.8100±0.10 | 0.7667±0.21 | 0.7667±0.16 | 0.7314±0.18 |
| | Transformer | 0.6373±0.06 | 0.6635±0.09 | 0.6065±0.08 | 0.5799±0.11 | 0.7490±0.19 | 0.7089±0.27 | 0.7566±0.18 | 0.7081±0.24 | 0.7593±0.02 | 0.6414±0.22 | 0.6658±0.14 | 0.6162±0.15 |
| Ours | BB-STT | **0.7817±0.05** | **0.7504±0.06** | **0.7504±0.06** | **0.7504±0.06** | **0.8118±0.17** | **0.7782±0.23** | **0.7782±0.23** | **0.7782±0.23** | **0.8723±0.14** | **0.8146±0.23** | **0.8146±0.23** | **0.8146±0.23** |

*Short-form video vs. mobile gaming* comparison yields the highest performance, with BB-STT achieving 87.23% accuracy, and 81.46% F1-score, representing the strongest result across all tasks. This result is counterintuitive given that both conditions involve active smartphone use but reflect the fundamentally different physiological profiles induced by passive reward-driven consumption and active cognitively demanding play. CNN-LSTM achieves 81.00% accuracy and 73.14% F1 on this comparison, confirming that the contrast is broadly learnable across architectures and reflects a robust underlying signal difference.

*Mobile gaming vs. baseline* viewing task also shows strong performance (with BB-STT, 81.18% accuracy, and 77.82% F1-score). Gaming produces the most pronounced physiological departure from any resting condition, reflected in strong performance across most model families. CNN-LSTM reaches 77.89% accuracy, and EEGNet 76.72% accuracy, consistent with the pronounced physiological contrast between active engagement and calm monitor viewing.

*Baseline vs. short-form video* scrolling is the most challenging pairwise task, with BB-STT achieving 78.17% accuracy, and 75.04% F1-score. This reflects the subtle nature of the distinction, as both conditions involve passive visual consumption and their physiological differences are genuinely small. Notably, the variance across folds is lower for this comparison (*±*0.05) than for gaming comparisons (*±*0.14–0.17), indicating that while absolute performance is lower, the signal is more consistent across subjects.

### E. Modality Ablation: Contribution of Individual Physiological Signals

Table VII and Fig. 2 present systematic modality ablation results across all five tasks and eleven modality combinations, revealing which physiological signals carry the most discriminative information for each behavioral contrast.

**Fig. 1.**
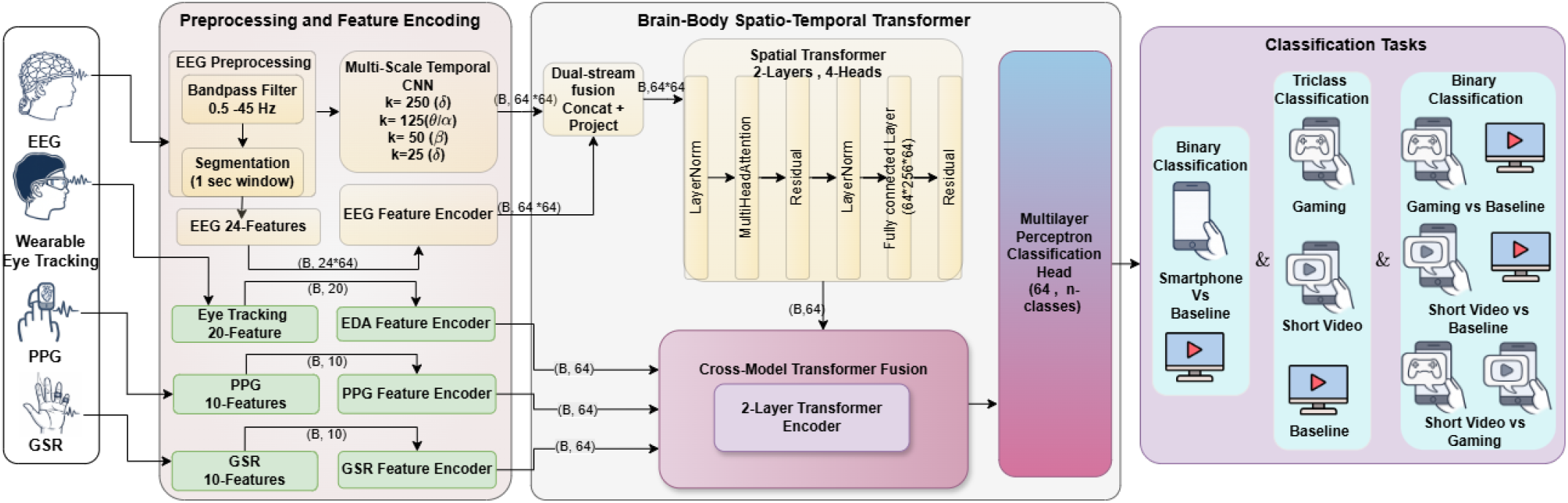
Overview of the proposed Brain-Body Spatio-Temporal Transformer (BB-STT) pipeline for multimodal affective-state decoding. Multimodal raw data (64-ch EEG, EDA, PPG, ET) is preprocessed and affective-relevant features are extracted for all modalities. The EEG encoder combines a multi-scale temporal CNN operating on raw EEG (64 ×1000) with a feature MLP operating on pre-extracted features (64× 24); both streams are concatenated and projected to produce per-channel embeddings (64× 64). A spatial transformer with a learnable [CLS] token models inter-electrode relationships. MLP encoders project peripheral affective features to 64-d embeddings, which are fused with the EEG representation via cross-modal attention. An MLP head produces task-specific affective-state predictions.

**Fig. 2.**
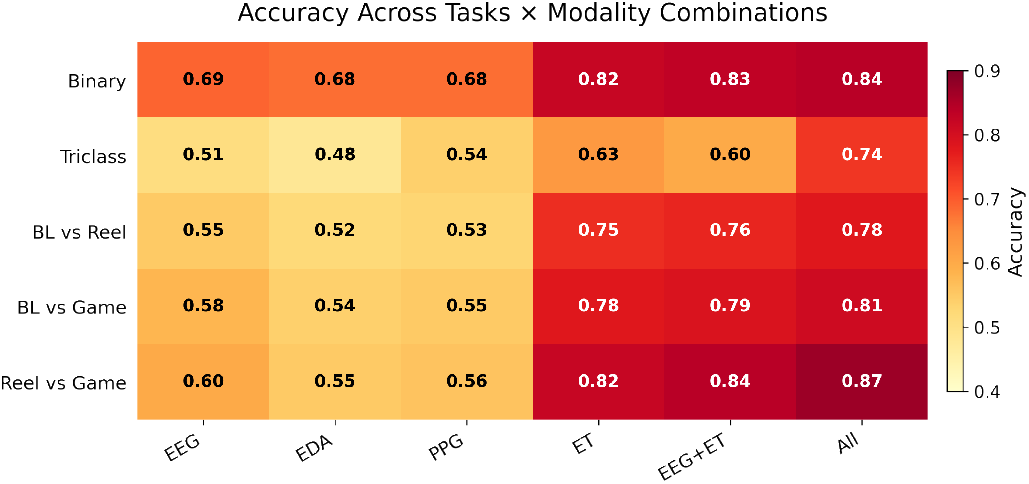
Affective-state decoding accuracy heatmap across five tasks and six modality configurations. ET consistently drives the largest performance gains.

Among single modalities, ET consistently provides the strongest performance across every task: 82.00% binary accuracy (F1: 76.84%), 62.73% three-class accuracy (F1: 52.44%), 79% on BL vs. SV, 82% on BL vs. Game, and 77% on SV vs. Game. ET outperforms EEG (68.54% binary, 51.39% three-class), PPG (67.80% binary, 53.96% three-class), and EDA (67.57% binary, 48.46% three-class) by substantial margins, reflecting the strong behavioral signature that gaze dynamics carry during smartphone interaction. The EEG and ET combination achieves 83.17% binary accuracy (F1: 78.66%) and 59.96% three-class accuracy (F1: 51.33%). The binary task shows a +1.17% gain over ET alone, indicating complementary neural contributions. EDA and PPG contribute selectively rather than uniformly across tasks. EEG with EDA underperforms EEG alone on binary accuracy (67.34% vs. 68.54%), indicating limited reliability at short temporal scales. PPG shows its strongest relative contribution in the SV vs. Game comparison, consistent with sensitivity to arousal differences. The full four-modality combination achieves the best overall performance on both binary (83.51%, F1: 78.35%) and three-class (74.13%, F1: 69.57%) tasks, confirming that each modality contributes non-redundant information when integrated through the cross-modal attention mechanism, even where individual pairwise combinations show diminishing returns.

### F. Architectural Ablation: Contribution of Each Component

Table VI presents the architectural ablation, isolating the contribution of the multi-scale temporal encoder, spatial transformer, and fusion mechanism across binary and three-class tasks.

**TABLE VI.** Architecture ablation study (5-fold cv). showing the effect of each component on binary and three-class affective-state decoding tasks.

| Configuration | Binary |  | Three-class |  |
| --- | --- | --- | --- | --- |
|  | Acc | F1 | Acc | F1 |
| *Temporal Encoder* |  |  |  |  |
| Single-scale (k=50 only) | 0.8265 | 0.7725 | 0.5814 | 0.4837 |
| Multi-scale (4 branches) | **0.8351** | **0.7835** | **0.7413** | **0.6957** |
| *Spatial Encoder* |  |  |  |  |
| Mean pooling (no transformer) | 0.8135 | 0.7437 | 0.6156 | 0.4862 |
| Learned GCN (ST-GNN style) | 0.8286 | 0.7754 | 0.5718 | 0.4702 |
| Spatial Transformer + CLS | **0.8351** | **0.7835** | **0.7413** | **0.6957** |
| *Fusion Mechanism* |  |  |  |  |
| No fusion (EEG only) | 0.6724 | 0.4781 | 0.4394 | 0.3439 |
| Concatenation | 0.8356 | 0.7893 | 0.5739 | 0.4328 |
| Gated fusion | 0.8282 | 0.7775 | 0.5858 | 0.4776 |
| Cross-attention | **0.8351** | **0.7835** | **0.7413** | **0.6957** |
| **BB-STT (Full)** | **0.8351** | **0.7835** | **0.7413** | **0.6957** |

**TABLE VII.** Modality ablation study (5-fold cv). showing the impact of each physiological signal on affective-state decoding.

| Modalities | Binary |  | Three-class |  |
| --- | --- | --- | --- | --- |
|  | Acc | F1 | Acc | F1 |
| EEG only | 0.6854 | 0.4565 | 0.5139 | 0.3778 |
| EDA only | 0.6757 | 0.4508 | 0.4846 | 0.3469 |
| PPG only | 0.6780 | 0.4773 | 0.5396 | 0.3938 |
| ET only | 0.8200 | 0.7684 | 0.6273 | 0.5244 |
| EEG + EDA | 0.6734 | 0.4230 | 0.4729 | 0.3651 |
| EEG + PPG | 0.6873 | 0.4859 | 0.5086 | 0.3621 |
| EEG + ET | 0.8317 | 0.7866 | 0.5996 | 0.5133 |
| EEG + EDA + PPG | 0.6702 | 0.4257 | 0.4923 | 0.3379 |
| EEG + EDA + ET | 0.8291 | 0.7772 | 0.5930 | 0.5508 |
| EEG + PPG + ET | 0.8259 | 0.7730 | 0.6311 | 0.5170 |
| **All (EEG+EDA+PPG+ET)** | **0.8351** | **0.7835** | **0.7413** | **0.6957** |

The temporal encoder comparison shows that replacing four parallel multi-scale branches with a single-scale design (k=50 only) reduces binary accuracy from 83.51% to 82.65% (F1: 77.25%) and three-class accuracy from 74.13% to 58.14% (F1: 48.37%). The three-class degradation of 15.99 percentage points is the largest single-component drop observed in the ablation, highlighting the importance of capturing multi-frequency dynamics for fine-grained discrimination.

The spatial encoder comparison reveals that mean pooling reduces three-class accuracy to 61.56% (F1: 48.62%) and that a learned GCN reduces it further to 57.18% (F1: 47.02%), despite the GCN improving binary accuracy over mean pooling (82.86% vs. 81.35%). The spatial transformer with CLS token achieves the best performance on both tasks, indicating the advantage of learning inter-electrode relationships through self-attention.

The fusion mechanism comparison provides the clearest architectural insight. Removing multimodal fusion entirely (EEG only) reduces binary accuracy to 67.24% (F1: 47.81%) and three-class accuracy to 43.94% (F1: 34.39%). Simple concatenation substantially recovers binary performance (83.56%, F1: 78.93%) but yields only 57.39% three-class accuracy (F1: 43.28%), indicating that naive fusion cannot resolve the subtler SV vs. gaming distinction. Gated fusion improves three-class performance modestly to 58.58% (F1: 47.76%), while cross-modal attention achieves 74.13% (F1: 69.57%), a 16.74-point improvement over concatenation on the three-class task. This substantial gain on the most challenging task highlights the importance of cross-attention, where selective reweighting of peripheral signals relative to EEG enables effective discrimination between interaction types with similar behavioral characteristics.

### G. Interpretability Analysis: SHAP Values, Feature Importance, and Spatial Attention

To understand how BB-STT resolves each discrimination, we combine permutation feature importance (ΔAccuracy), SHAP values (Fig. 3), and spatial transformer attention topographies maps (Fig. 4). Features in the top-right quadrant are individually irreplaceable and consistently influential, whereas those with high SHAP but moderate ΔAccuracy indicate redundancy with correlated features.

**Fig. 3.**
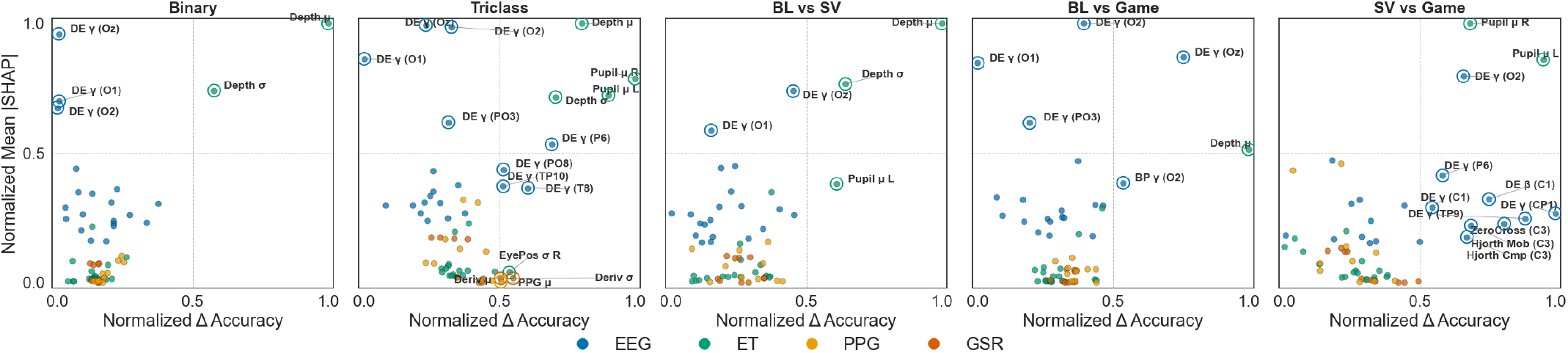
SHAP-permutation importance scatter plots for all five classification tasks. Features are plotted by mean —SHAP— (y-axis) against permutation-based accuracy drop ΔAccuracy (x-axis), colored by modality (green: ET, blue: EEG, orange: PPG, pink: GSR).

**Fig. 4.**
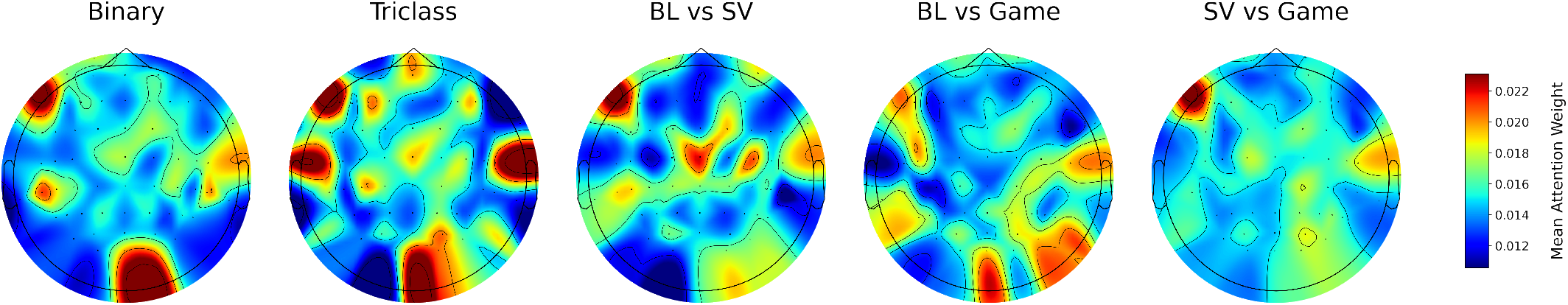
Topographic scalp maps of mean spatial transformer attention weights across 64 EEG channels for all classification tasks. Warmer colors indicate higher attention weights. Each map illustrates the relative importance of different brain regions for the corresponding discriminative task.

Across binary, BL vs. SV, and triclass tasks, gaze depth mean (Depth *µ*) occupies the top-right quadrant on both axes, establishing it as the primary discriminator for smartphone vs. non-smartphone engagement. The mechanism is geometric: smartphone use repositions the visual focal plane from a fixed-distance monitor to a handheld device at varying proximity, a signal no EEG, PPG, or GSR feature substitutes for. The binary and BL vs. SV spatial attention maps show elevated occipital and prefrontal electrode weighting, consistent with the visual system’s response to altered screen depth and frontal engagement differences between active device use and passive rest.

Differential entropy *γ* at occipital and parieto-occipital electrodes (Oz, O1, O2, PO3, P6) consistently appears in the high-SHAP/moderate-ΔAccuracy region across all five tasks, reflecting sustained visual cortical engagement that varies across all three behavioral conditions. A prominent occipital hotspot is present in binary, triclass, BL vs. SV, and BL vs. Game attention maps. For BL vs. Game specifically, occipital gamma achieves the highest SHAP values in the paper (ΔAcc up to +0.260 at electrode O2), with ET features entirely absent from the top rankings, confirming that gaming vs. baseline is a neural discrimination driven by sustained visuomotor tracking. The BL vs. Game attention map is correspondingly the most spatially diffuse, reflecting gaming’s simultaneous recruitment of visual, motor, and executive systems.

For SV vs. Game, the discriminative profile shifts fundamentally: bilateral pupil diameter (Pupil *µ* R and L) becomes the only ET feature in the top-right quadrant, reflecting differences in autonomic arousal. Central EEG features (differential entropy *γ* and *β* at C1, CP1, TP9; Hjorth measures at C3) enter the importance profile for the first time, capturing motor and executive engagement differences. The corresponding attention maps show more distributed weighting, suggesting the absence of a single dominant region (Fig. 4).

## VI. Discussion

### A. Smartphone Engagement as a Physiologically Coherent and Generalizable State

The binary result (83.51% accuracy, 78.35% F1) demonstrates that smartphone engagement produces a consistent physiological signature distinguishable from passive digital viewing regardless of smartphone interaction type. The LOSO evaluation strengthens this finding. LOSO accuracy (84.32%) marginally exceeds 5-fold CV (83.51%), demonstrating that BB-STT learns population-level signatures rather than subject-specific patterns. This generalization is explained by the interpretability finding that primary binary discriminators, ET gaze depth features, are geometric consequences of handheld device interaction shared across individuals. The selective robustness of gaming subjects in LOSO (sub-11: 92.44%, sub-07: 91.33%) further confirms that active gaming induces a strong and individually consistent engagement profile, while passive scrolling produces more individually modulated responses.

These findings align with Ward et al. [34], who demonstrated that smartphone presence alone reduces available cognitive capacity, and with Ciman and Wac [9], who showed that smartphone interaction gestures encode sufficient information for stress assessment. The present results extend these observations from behavioral performance measures and gesture-level signals to direct multimodal physiological decoding, establishing that the physiological fingerprint of smartphone engagement is sufficiently consistent across individuals and interaction types to support accurate cross-subject classification.

### B. Resolving Passive Consumption from Active Smartphone Engagement

The most theoretically significant classification finding is that short-form video scrolling and mobile gaming produce physiologically distinguishable states classifiable with 87.23% accuracy. The discrimination is not between smartphone engagement and non-smartphone engagement, but between qualitatively different forms of engagement that manifest differently across the full physiological hierarchy from cortex to autonomic nervous system.

For BL vs. game comparison, the differential entropy of the gamma band at occipital electrodes achieves the highest SHAP values in the paper (ΔAcc up to +0.260), with ET features entirely absent from the top rankings. This indicates that distinction is primarily neural, driven by sustained occipital gamma activity associated with visuomotor processing and attentional demands during gameplay, consistent with Hafeez et al. [6]. For SV vs. Game, the discriminative signal shifts to bilateral pupil dilation, the only task where pupillometry leads both SHAP and ΔAccuracy, reflecting the arousal differential between passive consumption and active cognitively demanding gaming. Yan et al. [2] and Pathak et al. [36] independently documented the diminished executive engagement and hedonic reward-seeking characteristic of short-form video consumption, consistent with the lower arousal reflected in the pupillometric distinction here. Central EEG features appearing uniquely in the SV vs. game profile further confirm that motor cortical dynamics distinguish active from passive smartphone engagement at the neural level.

### C. The Challenge of Passive-to-Passive Discrimination

BL vs. SV (78.17% accuracy, 75.04% F1) is the most challenging comparison, as both conditions involve passive visual consumption. The lower fold variance (*±*0.05 vs. *±*0.14–0.17 for gaming comparisons) indicates a weak but consistent signal. Gaze depth mean and standard deviation again dominate both axes for BL vs SV, confirming that the primary discriminator remains the geometric repositioning from stationary to handheld screen rather than any neural or autonomic engagement difference. The left-lateralized frontal-temporal spatial attention pattern for this comparison is consistent with the language and narrative processing associated with video content consumption. Scrolling involves reading captions and processing rapidly changing audiovisual content that stationary baseline viewing does not impose. The alpha band features at electrodes CP1 and P1 that contribute selectively to BL vs SV (ΔAcc ≈ +0.049–0.055) are consistent with posterior alpha desynchronization during short-form video consumption, reflecting increased visual cortical engagement, a pattern documented by Suhail et al. [51] as a neural correlate of social media engagement. ET alone achieves 79% on this comparison, only marginally below the full system, suggesting that the behavioral-geometric signal of handheld device use is the primary available discriminator and that EEG adds limited value for passive-to-passive discrimination.

### D. The Eye-Tracking Dominance Finding and Its Practical Implications

The modality ablation reveals that ET alone (82.00% binary accuracy) nearly matches the full BB-STT system (83.51%), which is consistent with SHAP analysis indicating that gaze depth features dominate across tasks involving baseline comparisons. Two interpretations of this finding are worth distinguishing. From a methodological perspective, ET may be capturing behavioral variance, the visible difference in gaze behavior between scrolling, gaming, and passive viewing, that is strong enough to drive most of the binary classification performance independently. From a scientific perspective, if gaze depth genuinely encodes the engagement distinction between device-interaction contexts, then ET is not merely a convenient proxy but a direct window into the attentional and spatial engagement processes that define each behavioral state [11]. The ablation and SHAP analyses highlight the complementary and task-specific role of EEG. While combining EEG with ET yields only modest gains in some settings, the overall pattern is informative. EEG contributes most precisely where behavioral-geometric differences are smallest, such as in the SV vs Game comparison where both conditions involve smartphone use and gaze depth loses its discriminative leverage. In contrast, when behavioral and geometric differences are more pronounced, gaze-based features dominate, and the contribution of EEG is reduced. This selective complementarity supports the multi-modal design. Neither ET nor EEG alone captures the full range of discriminative information across all tasks. Their integration through cross-modal attention enables dynamic weighting of modalities, resulting in consistently improved performance across different behavioral contrasts.

### E. Cross-Modal Attention as the Critical Architectural Mechanism

Cross-modal attention improves three-class accuracy by 16.74 points over concatenation, the largest gain in the study, while concatenation performs comparably on binary (83.56%). This divergence reflects the task structure: binary discrimination is dominated by ET gaze depth regardless of context, making fixed-weight fusion adequate, while three-class discrimination requires dynamic reweighting as the relative informativeness of EEG, ET, and pupillometry shifts across the three pairwise boundaries simultaneously. This is consistent with Ding et al. [25] and Zhang et al. [26], and extends their findings to behavioral classification, demonstrating that the cross-attention advantage concentrates at the hardest classification boundary. The 15.99-point three-class drop from multi-scale to single-scale temporal encoding reflects the triclass task’s requirement to simultaneously capture delta/theta dynamics associated with reward processing and beta/gamma dynamics associated with visuomotor engagement. This multi-scale requirement is consistent with the SHAP finding that the triclass task recruits EEG features across delta, theta, alpha, and gamma bands simultaneously, a multi-band engagement profile that a single temporal scale cannot adequately capture.

### F. A Two-Stage Physiological Hierarchy: Implications for System Design

The interpretability analysis reveals a two-stage physiological hierarchy that provides a mechanistic account of the full classification system and has direct implications for practical deployment. The first stage, separating smartphone from non-smartphone engagement, is solved primarily by ET gaze depth. A geometric signal reflecting the fundamental spatial repositioning from stationary to handheld screen use, reinforced by occipital EEG gamma as a stable visual cortical engagement index. The second stage, resolving which type of smartphone engagement is occurring, requires EEG with task-specific frequency bands and electrode clusters: broad lateral occipital and central-temporal gamma for gaming vs. baseline, frontal alpha and delta for scrolling vs. baseline, and central EEG complexity combined with bilateral pupil dilation for the within-smartphone SV vs. gaming discrimination. The practical implication is a tiered sensing architecture: a lightweight ET-only system for coarse context detection, augmented by EEG when fine-grained behavioral discrimination is required. For digital wellbeing applications specifically, Islambouli et al. [33] showed that purposeless no-goal smartphone sessions are associated with regret and reduced wellbeing, and Nguyen et al. [3] meta-analytically confirmed negative associations between heavy short-form video use and attention control. The ability to passively infer whether a user is in a high-arousal active gaming state or a low-arousal habituated scrolling state, without any explicit user input, opens a path toward objectively characterizing the quality of digital engagement rather than its mere quantity. The present work establishes the physiological feasibility of this discrimination and identifies an efficient multimodal sensing strategy to achieve it.

### G. Limitations and Future Directions

The dataset comprises 21 participants with a 2:1 SV-to-gaming imbalance from self-selection, producing higher variance in gaming-related task estimates. Future work should recruit balanced groups across game genres, difficulty levels, and content platforms. Additionally, short-form video and gaming content varied across participants, so the classified signatures may partially reflect content-specific rather than interaction-type-specific differences; future designs should control content within groups or include content type as a covariate. All data were recorded in a single session, precluding assessment of test-retest reliability, habituation effects, and time-of-day confounds; multi-session longitudinal designs are needed to evaluate temporal stability. Finally, generalization across recording sessions, device types, and demographic groups remains to be established; domain adaptation and larger naturalistic datasets represent the primary directions for future work.

## VII. Conclusion

This study demonstrates that distinct smartphone interactions, such as passive short-form video scrolling and active mobile gaming, produce distinguishable multimodal physiological signatures. To decode these states, we introduce the Brain-Body Spatio-Temporal Transformer (BB-STT), a unified multimodal framework that integrates neural and peripheral physiological signals through cross-modal attention. A two-stage physiological hierarchy further emerges, in which gaze depth supports device-context separation, while combined pupillometric and EEG features enable fine-grained discrimination between smartphone interaction types.

These findings suggest that smartphone usage may not be adequately characterized as a homogeneous activity but rather reflects physiologically diverse patterns. They support the feasibility of real-time, passive assessment of smartphone engagement quality in naturalistic settings. Overall, this work highlights the potential of multimodal physiological signals for decoding interaction-specific engagement and enabling future context-aware, human-centered digital systems.

